# Nonlinear dendritic integration supports Up-Down states in single neurons

**DOI:** 10.1101/2024.09.05.611249

**Authors:** Alessio Quaresima, Hartmut Fitz, Peter Hagoort, Renato Duarte

## Abstract

Changes in the activity profile of cortical neurons are due to effects at the scale of local and long-range networks. Accordingly, abrupt transitions in the state of cortical neurons—a phenomenon known as Up-Down states—have been attributed to variation in the activity of afferent neurons. However, cellular physiology and morphology may also play a role in causing Up-Down states. This study examines the impact of dendritic nonlinearities, particularly those mediated by voltage-dependent NMDA receptors, on the response of cortical neurons to balanced excitatory/inhibitory synaptic inputs. Using a neuron model with two segregated dendritic compartments, we compared cells with and without dendritic nonlinearities. NMDA receptors boosted somatic firing in the balanced condition and increased the correlation between membrane potentials across the compartments of the neuron model. Dendritic nonlinearities elicited strong bimodality in the distribution of the somatic potential when the cell was driven with cortical-like input. Moreover, dendritic nonlinearities could detect small input fluctuations and lead to Up-Down states whose statistics and dynamics closely resemble electrophysiological data. Up-Down states also occurred in recurrent networks with oscillatory firing activity, as in anaesthetized animal models, when dendritic NMDA receptors were partially disabled. These findings suggest that there is a dissociation between cellular and network-level features that could both contribute to the emergence of Up-Down states. Our study highlights the complex interplay between dendritic integration and activity-driven dynamics in the origin of cortical bistability.

**Significance statement:** In several physiological states, such as sleep or quiet wakefulness, the membrane of cortical cells shows characteristic bistability. Cells are either fully depolarized and ready to spike, or in a silent, hyperpolarized state. This dynamics, known as Up-Down states, has often been attributed to changes in network activity. However, whether cell-specific properties, such as dendritic nonlinearities, play a role in neuronal bistability remains unclear. This study uses a dendritic model of a pyramidal cell and shows that the presence of NMDA receptors drives the Up-Down states in response to small fluctuations in network activity. Thus, single cells can enter the Up-Down state dynamics independently of the ongoing network activity.

## Introduction

Cortical neurons exhibit diverse activity patterns that vary with the behavioral and internal states of an animal (Flavell *et al*., 2022; Greene *et al*., 2023). A common view is that these patterns are primarily regulated by network-level phenomena at local and global scales (Harris & Thiele, 2011; Poulet & Crochet, 2019). Network activity dominates the cellular state by activating excitatory and inhibitory synapses, which are fine-tuned by plasticity (Shu *et al*., 2003; Higley & Contreras, 2006; Zhou & Yu, 2018). However, recent experimental and computational results have highlighted how cellular features, such as dendritic arborization and synaptic organization, influence neuronal responses to network inputs (Iascone *et al*., 2020; Otor *et al*., 2022; Dembrow & Spain, 2022; Larkum, 2022; Gidon *et al*., 2020; Lafourcade *et al*., 2022) and account for variability in the expression of cortical states across layers and cell types (Poulet & Crochet, 2019; Tukker *et al*., 2020).

Due to synaptic bombardment, the membrane conductance increases, leading to the so-called high-conductance state (Destexhe *et al*., 2003; Okun & Lampl, 2008; Zhou & Yu, 2018; Ebsch & Rosenbaum, 2018). This state has been measured *in vivo* and is considered the operating point of cortical cells in awake animals (Destexhe *et al*., 2007; Poulet & Crochet, 2019; McGinley *et al*., 2015). However, it is only one aspect of the more complex cortical dynamics. In certain physiological states of the animal, such as anesthesia, sleep, or quiet wakefulness, the somatic membrane of cortical neurons has a bimodal distribution and fluctuates around two metastable states known as Up-Down states (UDS, Cowan & Wilson, 1994; Wilson & Kawaguchi, 1996; Steriade *et al*., 2001). In the Up state, the cell is depolarized, and somatic firing is driven by fluctuations in the balance of transmembrane excitatory and inhibitory currents, similar to the high-conductance state (Destexhe *et al*., 2003). The Down state corresponds to a hyperpolarized neuron with a membrane potential close to rest. The prevalence of Up and Down states, as well as their duration and frequency, has been associated with the switching of local cortical circuits from inactive to active processing (Anderson *et al*., 2000; Luczak *et al*., 2007; Jercog *et al*., 2017; Tukker *et al*., 2020).

Network simulations and analytical insights have shown that two conditions are necessary for the emergence of Up-Down dynamics, balanced excitation/inhibition and strong excitatory feedback (Sanchez-Vives & McCormick, 2000; Compte *et al*., 2003; Droste & Lindner, 2017; Holcman & Tsodyks, 2006; Destexhe, 2009; Tartaglia & Brunel, 2017). However, network models often require fine-tuning of connectivity and other parameters to avoid pathological behavior and elicit the desired membrane dynamics (Renart *et al*., 2007; Maksimov *et al*., 2018). Experimental studies have proposed a different physiological explanation for UDS, emphasizing the relationship between Up states and the onset of plateau potentials in dendrites (Milojkovic *et al*., 2005; Antic *et al*., 2010; Oikonomou *et al*., 2014). In this view, although network activity triggers the Up state, its bistability is shaped by voltage-dependent NMDA receptors (NMDARs) and other regenerative dendritic events (Larkum, 2013; Antic *et al*., 2010; Papoutsi *et al*., 2014). However, recent experimental findings have shown intact UDS despite NMDAR-blockage (Palmer *et al*., 2014; Smith *et al*., 2013; Chen *et al*., 2012), suggesting that local dendritic balance, rather than dendritic nonlinearities, may be required for the emergence of UDS (Larkum, 2022).

Conflicting evidence also derives from other modeling studies. Benucci *et al*. (2004) showed that including voltage-dependent receptors in models with realistic morphology leads to UDS in response to pairwise correlations in the input, but not for simplified dendritic structure. In contrast, Papoutsi *et al*. (2014) showed that the expression of NMDARs in neurons with a single dendritic compartment is sufficient for the onset and maintenance of Up states in small networks. Crucially, these studies did not implement local dendritic balance, leaving two key questions unanswered: first, are fluctuations in the dendritic E/I balance sufficient to induce UDS in simplified models? And secondly, how does the nonlinearity of NMDARs interact with the local E/I-balanced state?

This article investigates the relationship between dendritic E/I balance and Up-Down states. We propose a unified description of these phenomena using a neuron model that reproduces key features of nonlinear dendritic integration (Quaresima *et al*., 2022). After validating the physiological response of the model to a barrage of E/I-balanced synaptic inputs, we study its dynamics with and without NMDARs. We demonstrate that NMDARs strongly affect firing activity and the cell’s internal state in that they enhance sensitivity to correlations in the input. We show that NMDARs support the occurrence of cellular UDS under awake-like cortical activity. When NMDARs are blocked, UDS requires network oscillations, as observed in anesthetized animals. Therefore, we advance the hypothesis that network-level Up-Down states are sufficient but not necessary for bimodality in the distribution of cellular membrane potentials.

## Materials and Methods

### Dendritic neuron model

We investigate the balanced state and Up-Down dynamics in a model of dendritic integration which consists of three computational elements, or compartments (Quaresima *et al*., 2022). The neuron has an axosomatic compartment, representing the soma and perisomatic segments, and two electrotonically segregated passive dendritic compartments that are coupled to the soma with membrane timescales determined by their dendritic lengths (in the range of 100 µm to 500 µm). The neuron model was characterized for two sets of parameters from human and mouse single-cell experiments (Quaresima *et al*., 2022). In the present study, we used human parameters for dendritic and synaptic physiology, which were shown to exhibit stronger nonlinear NMDAR-mediated effects.

### Axosomatic and dendritic compartments

The axosomatic compartment is formalized as an adaptive exponential integrate-and-fire model (Eqs. 1, 2, and 3; Brette & Gerstner, 2005), parameterized for the adaptive firing regime (Table 1). Dendritic compartments were approximated as conductive cylinders whose voltage was governed by a passive membrane-patch equation similar to the soma, but lacking mechanisms for spike generation and intrinsic adaptation (Eq. 3). The three compartments are described by the following system of differential equations. For the somatic compartment:

**Table 1.** Axosomatic compartment. Parameters for the axosomatic compartment which is modeled as an adaptive exponential neuron. (Brette & Gerstner, 2005).

| Symbol | Description | Value | Unit |
| --- | --- | --- | --- |
| $g_L$ | Membrane leak conductance | 40 | nS |
| $C_m$ | Membrane capacitance | 281 | pF |
| $V_r$ | Resting membrane potential | -70.6 | mV |
| $V_T$ | Threshold potential | -50.4 | mV |
| $\theta$ | Spike onset threshold | 0 | mV |
| $u_r$ | Reset potential | -55 | mV |
| $\Delta_T$ | Slope factor | 2 | mV |
| $\tau_w$ | Spike-triggered adaptation time scale | 144 | ms |
| $a$ | Sub-threshold adaptation conductance | 4 | nS |
| $b$ | Spike-triggered adaptation increment | 80.5 | pA |
| $t_{up}$ | Spike width (soma clamped at 20 mV) | 1 | ms |
| $t_{ref}$ | Absolute refractory period | 2 | ms |

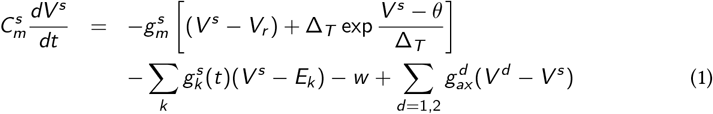

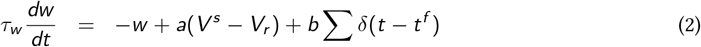

When the somatic voltage *V* ^*s*^ crosses a firing threshold *θ, V* ^*s*^ is set to the reset potential *u*_*r*_, *θ* is increased by 10 mV and decays back to baseline with time constant of 30 ms. The two adaptation mechanisms (*θ* and *w*) model spike afterhyperpolarization on short and long timescales. For the two dendritic compartments (*d* = 1, 2):

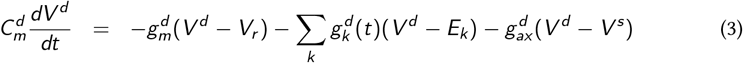

In this notation, the superscripts *s* and *d* denote the soma and dendrites, respectively. The parameters *C*_*m*_, *g*_*m*_, and *g*_*ax*_ refer to the capacitance of the membrane patch, the leak conductance, and the axial conductance. The values are computed using the cable equation:

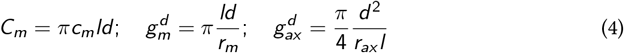

where *l* and *d* refer to the length and diameter of the dendritic cylinder. The physiological parameters that were used are those obtained by Eyal *et al*. (2016) in human cells, i.e., *r*_*m*_ = 39 kΩ cm^2^ (membrane resistance), *r*_*ax*_ = 200 Ω cm (intracellular resistance), *c*_*m*_ = 0.5 µF/cm^2^ (membrane capacitance). The current flow from the dendrites is dromic (depolarizing current to the soma, 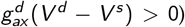, except when the neuron emits a spike. Parameters for the somatic and dendritic compartments are shown in Table 1 (soma) and Table 2 (dendrites). They are identical to Quaresima *et al*. (2022), except for the reset potential *u*_*r*_ which was set to −55 mV, following Bono & Clopath (2017).

**Table 2.** Dendritic compartments. Parameters for proximal and distal dendritic compartments in the neuron model.

| Symbol | Description | Distal | Proximal | Unit |
| --- | --- | --- | --- | --- |
| $l$ | Dendritic length | 400 | 150 | $\mu\text{m}$ |
| $d$ | Dendritic diameter | 4 | 4 | $\mu\text{m}$ |
| $g_m$ | Leak conductance | 1.29 | 0.32 | nS |
| $\tau_d$ | Membrane time constant | 1.48 | 0.22 | ms |
| $g_{ax}$ | Axial conductance | 15.71 | 62.83 | nS |
| $C_m$ | Membrane capacitance | 25.13 | 6.28 | pF |

### Synaptic dynamics

We modeled excitatory glutamatergic transmission as mediated by fast AMPA and slow NMDA receptors, and inhibitory GABAergic transmission by fast GABA_A_and slow GABA_B_receptors. Receptors were modeled as conductances with double-exponential kinetics (Roth & van Rossum, 2009):

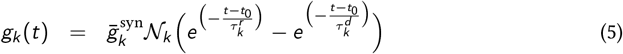

where *t*_0_ is the pre-synaptic spike time, *k* ∈ {AMPA, NMDA, GABA_A_, GABA_B_}, and 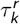 and 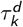 are the specific rise and decay time constants of each receptor (see Table 3). The maximal conductance of each receptor is given by 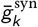 and scaled by a fixed normalization factor 𝒩_*k*_ :

**Table 3.**
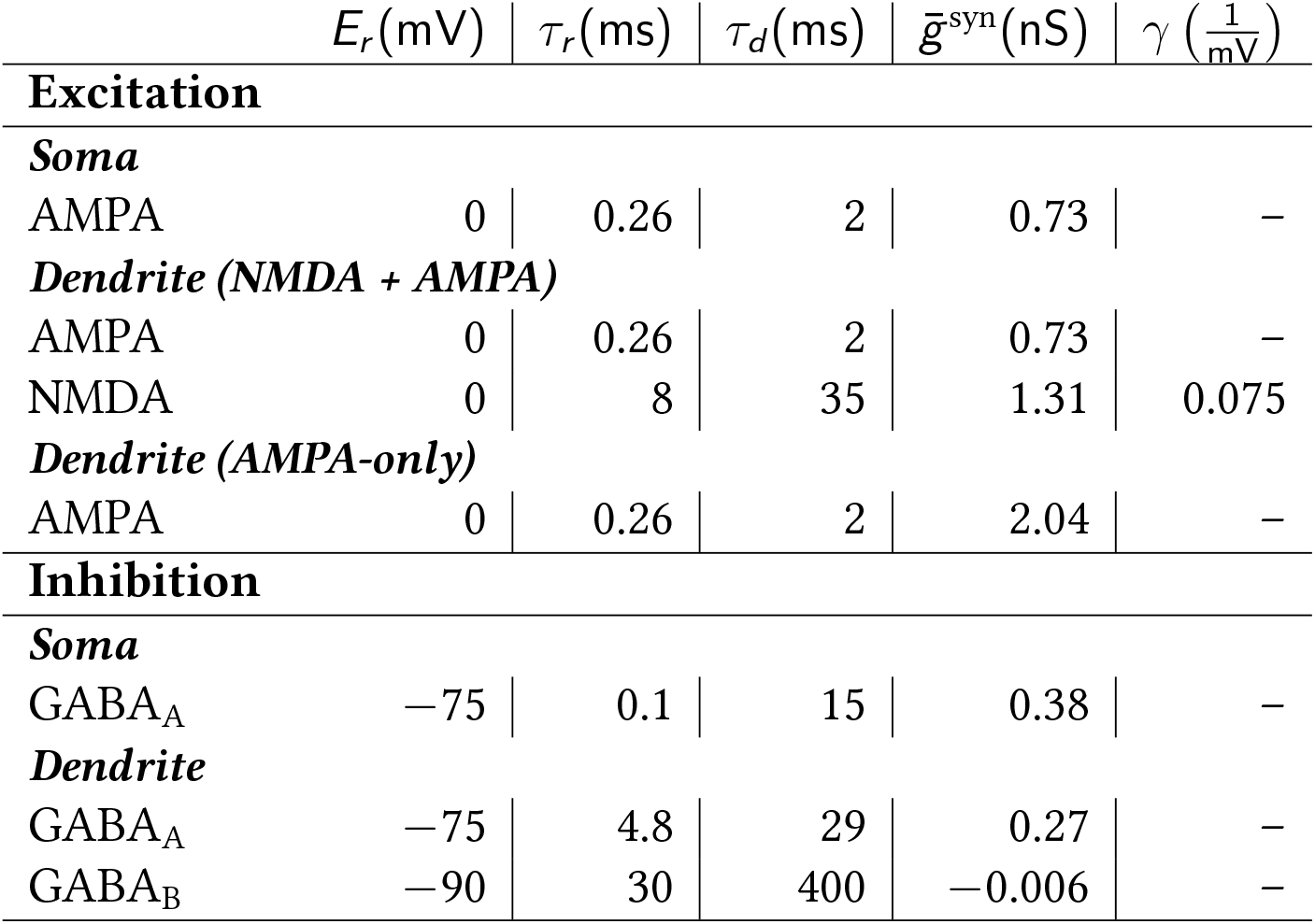
Synaptic parameters of the neuron model. *E*_*r*_ is the synaptic reversal potential, *τ*_*r*_ and *τ*_*d*_ are the conductance rise and decay time constants, 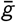 is the peak conductance, and *γ* determines the slope of the NMDA voltage-dependence.

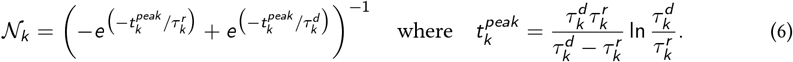

The conductance of the NMDA receptor was further scaled by a multiplicative voltage-gating mechanism *G* to capture its dependence on the dendritic membrane potential:

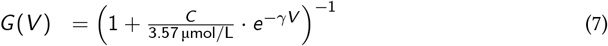

where *V* is the membrane voltage of the postsynaptic compartment and *γ* is a physiological parameter that controls the steepness of the voltage dependence. The extracellular concentration *C* of magnesium ions Mg^2+^ was fixed at 1 µmol*/*L. Equations and parameters for the NMDA receptor were based on Jahr & Stevens (1990).

To investigate the role of NMDARs in regulating the onset and duration of Up states, we compared synaptic configurations with and without NMDARs. In the latter condition (labeled AMPA-only), the AMPAR conductance was scaled such that its peak amplitude matched the sum of AMPA and NMDA receptor conductances in the condition with NMDARs. In this way, we could focus on two characteristics of NMDAR activation, its voltage-dependence and timescale. All synaptic parameters are shown in Table 3.

### Balance via modulation of the excitatory and inhibitory rate

Cortical neurons are exposed to intense, continuous synaptic bombardment from local and long-range afferents. During awake, active processing, excitatory and inhibitory input streams are balanced such that the soma of a post-synaptic neuron remains depolarized and fires sparsely at low rates (Harris & Thiele, 2011; Destexhe *et al*., 2003). We instantiated this condition by stimulating the neuron with Poisson-distributed excitatory and inhibitory spikes at high rates, mimicking inputs from thousands of afferents. Excitatory and inhibitory synaptic strengths were fixed throughout the study (Table 3), except when specified, but the input rates *ν*_*exc*_ and *ν*_*inh*_ were systematically varied (Fig. 1C) according to 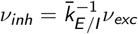, where 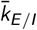 is determined such that the somatic membrane potential is in the balanced state (see below).

**Figure 1:**
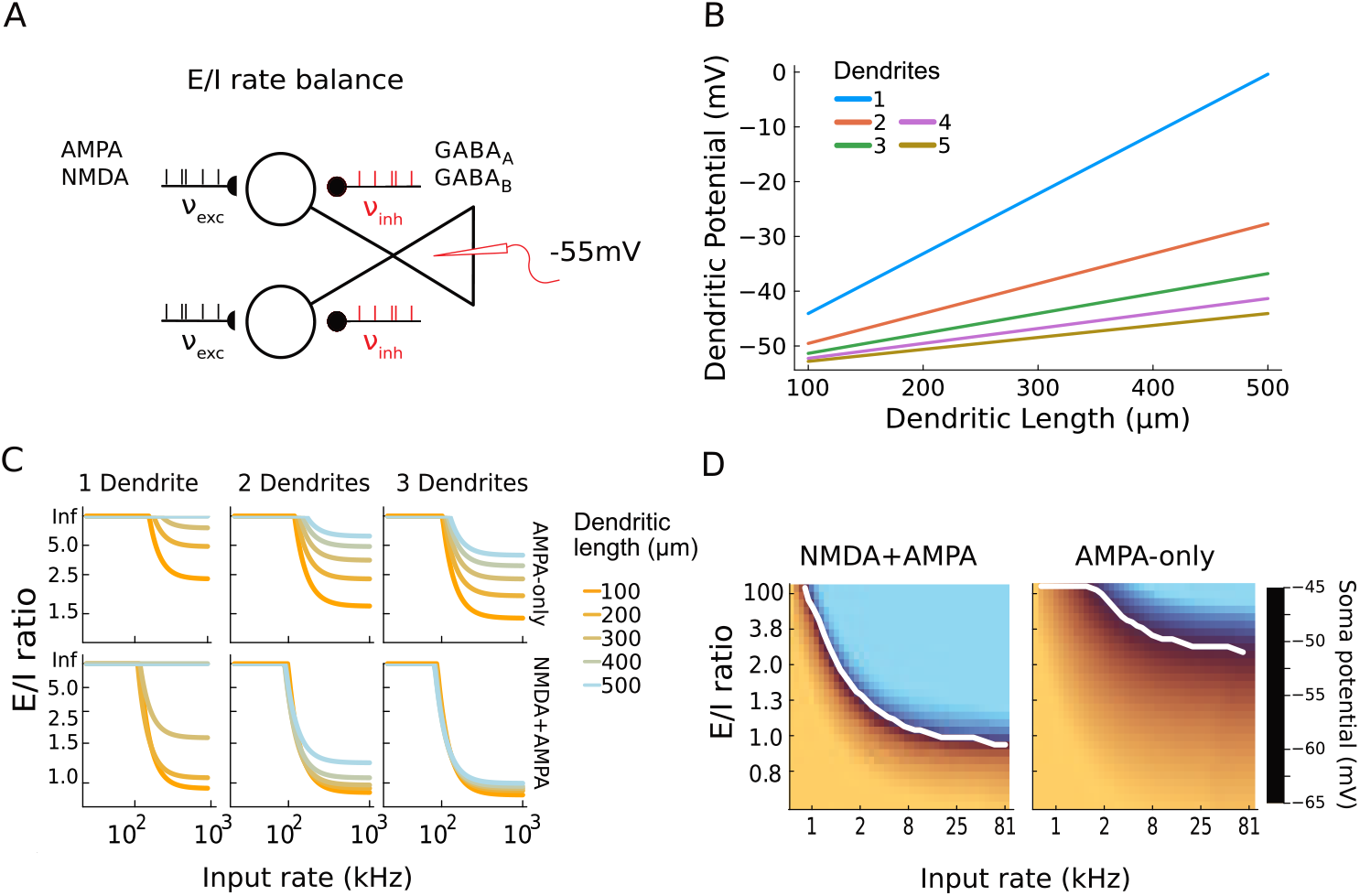
Excitation-inhibition balance in dendrites. **A**, Schematic of the protocol used to set the E/I balance. For a fixed excitatory input rate *ν*_*exc*_ on AMPA and NMDA receptors, we modulated the inhibitory firing rate *ν*_*inh*_ (red) onto GABA_A_ and GABA_B_ receptors until the membrane potential stabilized around − 55 mV. **B**, Dendritic membrane potential necessary to set the soma in the balanced condition depends on dendritic length and the number of dendritic compartments. **C**, The balanced E/I ratio was calculated analytically and depended on the excitatory input rate, dendritic length, number of dendritic compartments, and the receptor types that were expressed. **D**, Numerically determined 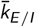. Panels show somatic membrane potential in neurons with dendrites of length 300 µm, with and without NMDARs. White lines indicate the target somatic potential (−55 mV). The 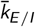 agree with the analytic approach.

The neuron’s membrane potential is balanced when the sum of leak and axial currents on the soma is zero. For a fixed leak conductance and a target somatic potential 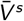, this condition is determined by the current that the dendritic compartments send to the soma which, in turn, depends on the ratio of excitation and inhibition on the dendrites (Fig. 1*A*). Knowing the dendritic length (*L*_*d*_) and the number of dendritic branches (*N*_*d*_), from Eq. 1 and Eq. 3 we can analytically determine the amplitude of the axial current required to reach this balanced state, and thus the dendritic membrane potential (Fig. 1*B*):

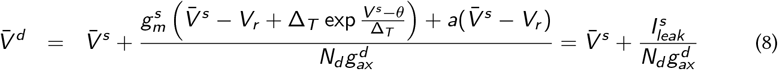

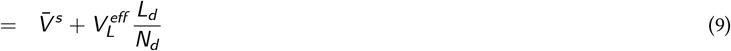

Note that, for simplicity, the axial conductance 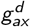 is expressed as a function of dendritic length (Eq. 4), with the remaining factor lumped into 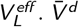 increases linearly with dendritic length and is proportional to the number of dendritic compartments. As shown in Fig. 1*B*, dendritic voltage remains within physiological bounds for dendritic lengths shorter than 500 µm, but it requires the dendrite to be fully depolarized if *N*_*d*_ = 1.

From Eq. 3 and Eq. 9, and assuming the neuron receives Poisson-distributed spike input with rates 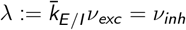, we determine the ratio between excitation and inhibition that achieves a stationary somatic potential, which we fix at 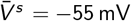. In this condition, the synaptic conductances can be replaced with their average values (following Campbell’s theorem, Kuhn *et al*., 2004):

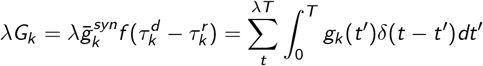

Thus, we obtain:

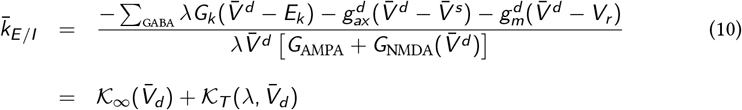

where

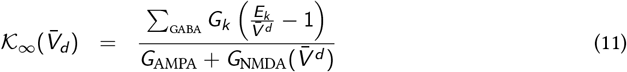

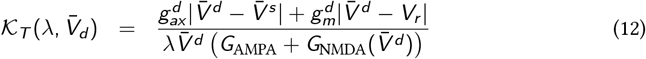

Note that, in Eq. 11 and Eq. 12, the reversal potentials for the glutamatergic receptors were removed because their values are *E*_AMPA_ = *E*_NMDA_ = 0 mV. From Eq. 9 and Eq. 10, we can determine the values of 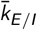 that establish balance as a function of dendritic length and input rate. For simplicity, we assume that the dendrites are symmetric, however the results can be generalized if the current flow from each dendritic compartment is assumed to be equal. First, 𝒦 _𝒯_ is strictly positive and, for high input rates, 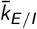 converges to 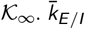 approaches the asymptotic value as the rate-induced excitatory conductance dominates the axial and leak conductances (high conductance state, Eq. 12). Secondly, the derivative 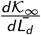 indicates that 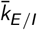 increases with dendritic length, but its dependence is attenuated by the voltage-dependent receptor. The analytically derived 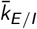 corresponding to different dendritic lengths, input rates, and synaptic conditions are shown in Fig. 1*C*.

To validate the analytical derivations, the 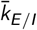 values were also computed numerically by simulating the neuron in the free membrane condition (no spike-threshold, *θ* → _∞_ in Eq. 1). We determined the average membrane potential for excitatory rates in the range of 0.5 kHz to 80 kHz and varied the ratio between excitatory and inhibitory inputs. For each excitatory rate, we selected the 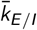 such that the average free membrane potential across 10 simulations was closest to the somatic target potential of − 55 mV (results shown in Fig. 1*D*). Additionally, we further validated this approach by considering an alternative definition of the balanced state, focused on supra-threshold activity and characterized by sparse, low-rate somatic firing at a fixed target rate of 5 Hz. This alternative yielded similar results for the value of 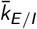 that is required to establish balance (data not shown).

### Experimental design and statistical analysis

#### Input fluctuations

It has been argued that Up-Down state transitions are due to fluctuations in the balance of E/I inputs (Benucci *et al*., 2004; Jercog *et al*., 2017). To study the role of E/I variations, we introduced controlled fluctuations in the firing rate *ν*_*exc*_ (*t*) of the excitatory inputs, following the method described in Vogels *et al*. (2011). We used a rectified Ornstein–Uhlenbeck process *z*_*t*_ with autocorrelation time *τ* = 50 ms which defines the instantaneous rate *r*_*t*_ of an inhomogeneous Poisson process, defined as follows:

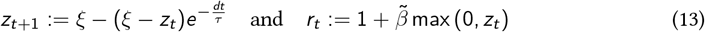

where *ξ* is a random variable in 𝒰_[*−* 0.5,0.5]_ and 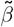 is a free parameter that scales the fluctuation size of the rectified process *r*_*t*_. We normalized the average of *ν*_*exc*_ (*t*) such that it was equal to the baseline input rate 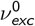 over the entire simulation interval. Thus, the instantaneous input rate of excitatory afferents is:

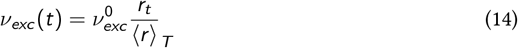

Drawing spikes from a Poisson distribution with rate *ν*_*exc*_ (*t*) yields a spike train with controlled statistics (mean rate and coefficient of variation of the inter-spike-intervals, CV_ISI_), as portrayed in Fig. 3*A*. We further introduce the parameter 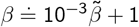, as a linear transformation of 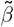, to ensure the CV_ISI_ of the generated spike train is in the range of 1 to 2, allowing us to use it as a biologically meaningful proxy for the fluctuation size of *ν*_*exc*_ (*t*).

#### Quantifying Up-Down states

The all-time-points histogram (APH) is a measure of the average time the membrane potential falls within a certain voltage range. It was first used in Wilson & Kawaguchi (1996) to characterize *in vivo* intracellular recordings from cortical and striatal spiny neurons. In the presence of Up-Down states, the APH shows a bimodal distribution of somatic membrane potentials. Values most frequently assumed by the membrane are either near the firing threshold (Up state) or close to the resting potential (Down state). We computed the APH for simulations of 10 s unless otherwise specified. The APH has a resolution of 1 mV.

Bimodality in the APH distribution was determined via Gaussian kernel density estimation (GKDE). Following Silverman (1981), we computed the GKDE for the APH as:

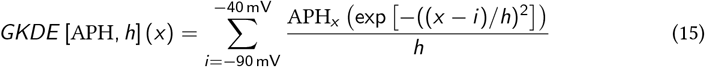

where APH_*x*_ is the magnitude of the histogram bin corresponding to the voltage *x*. The function *GKDE* [APH, *h*] (*x*) is a smooth curve that depends on the particular realization of the APH and the window size *h* used for the Gaussian convolution. It was demonstrated that, for a function obtained from Eq. 15, the number of local maxima of *GKDE* (*x*) is a monotonically decreasing function of *h* (Silverman, 1981). Thus, to estimate the bimodality of an APH, we computed the GKDE for windows in the range of 1 mV to 50 mV and counted the number of maxima. Because this number is monotonically decreasing, we can use the minimal window size 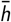 for which there is a unique maximum in the GKDE (unimodal distribution) as an index of the APH bimodality. The larger 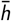 is, the stronger is the APH bimodality, indicating the presence of Up-Down states.

#### Network simulation

The network consisted of 800 excitatory cells, modeled in the same way as described above, and 200 inhibitory neurons modeled as soma-only leaky integrate-and-fire neurons. The parameters for inhibitory neurons were *g*_*m*_ = 5 nS (leak conductance), *θ* = −50 mV (spike threshold), *V*_*r*_ = −60 mV (resting potential), *u*_*r*_ = −60 mV (reset potential), and *C* = 100 pF (membrane capacitance). The network used two types of fast, double-exponential synapses with 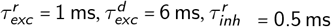, and 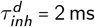 where *r* refers to the rise time and *d* to the decay time constant. The two populations were recurrently connected with probability 0.2 and synaptic weights of 2 pF, except for excitatory-to-inhibitory connections which had a strength of 0.5 pF. Within the inhibitory population, 35% of the cells targeted the somatic compartment of the excitatory population, mimicking parvalbumin-expressing cells, the remaining ones targeted the two dendrites with independent probability.

#### Numerical integration

Differential equations were integrated using Heun’s explicit method (Ascher & Petzold, 1998) with a step-size of 0.1 ms. Synaptic integration at high input rates (beyond 50 kHz) created numerical instability. When synaptic conductances became very large, sudden changes in the membrane potential (for example when the cell fires) lead to large incoming synaptic currents. To resolve this issue, we limited the derivative of the membrane potential by clipping the maximal current that flows through ionic channels at 1 nA.

#### Simulations

Simulations of single neurons were performed in Julia and the code is available on GitHub. Simulations of networks and neurons with several dendritic branches were carried out in a custom Julia library for spiking neural networks, JuliaSNN.

## Results

### E/I balance depends on receptor types and dendritic length

We investigated the impact of dendritic nonlinearities when cortical cells are under E/I-balanced synaptic bombardment. To this aim, we used a model of dendritic integration that has been shown to reproduce long-lasting dendritic depolarization known as plateau potentials, following NMDA spikes in proximal and distal dendrites (Quaresima *et al*., 2022). The NMDA receptors can be switched off to compare the dynamics of dendritic integration with and without voltage-dependent nonlinearity.

We first tested the model’s physiological validity in cortical-like regimes by exposing the dendrites to intense synaptic bombardment and measuring somatic activity. Dendrites govern the neuronal transfer function and somatic firing depends on input rate and dendritic geometry. However, for large input rates the neuron’s firing pattern is outside physiological ranges for most dendritic configurations. This issue arises from an imbalance of excitatory and inhibitory afferents in the dendritic compartments. We balanced E/I inputs such that the somatic membrane potential and firing rate resembled the high-conductance state (Destexhe *et al*., 2003). In line with experimental evidence and previous computational studies (Wilson, 2008; Kuhn *et al*., 2004), we defined E/I-balance in terms of the average membrane potential *V*_*s*_ of the somatic compartment. The balance was achieved by modulating the inhibitory rate such that *V*_*s*_ hovered below the threshold potential for spike generation (Fig. 1*A*, see details in Materials and Methods). We chose *V*_*s*_ = − 55 mV because it corresponds to the average potential of the Up state measured *in vivo* (Wilson & Kawaguchi, 1996) and was used in previous modeling studies (Kuhn *et al*., 2004). The somatic target voltage determines the dendritic potential that is required such that axial currents depolarize the soma up to *V*_*s*_. Neurons with longer dendrites require strongly depolarized dendrites and increasing the number of dendritic compartments relaxes the dependence on length (Eq. 9, Fig. 1*B*).

We obtained the excitatory/inhibitory ratio for the balanced condition analytically (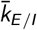, Eq. 10 and Fig. 1*C*) and verified that simulations lead to the same (analytic) 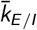 for models with and without NMDARs (Fig. 1*D*). The balance is geometry-dependent, with longer dendrites requiring stronger excitation (i.e., a larger 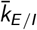) than shorter ones (Fig. 1*C*). However, voltage-dependent channels in the segregated compartments compensate for the lower axial conductance and reduce the difference between long and short dendrites. Models with NMDARs have a lower E/I ratio than models expressing only AMPARs because the opening of NMDARs causes long-lasting influx of depolarizing currents.

### NMDARs boost firing activity in the E/I-balanced state

To gain insights into how segregated dendrites and nonlinear integration due to NMDARs affect the neuronal dynamics in the balanced state, we compared the membrane potential of neuron models with and without voltage-dependent receptors when the model approaches the balance (input frequencies in the range 0.3 kHz to 5 kHz, Fig. 2*A*). The neuron reaches the E/I balance potential for sufficiently large input rates (dashed lines: NMDARs: 0.7 kHz, AMPARs: 1.5 kHz). While approaching the balanced state, the average dendritic potential depends on dendritic length. When NMDARs are present, the dendritic potentials split into two groups: those that enable E/I balance and those that do not (below and above the dashed line). The separatrix loosely matches the membrane potential that activates the voltage gate of NMDARs (∼− 50 mV). When excitatory input is sufficient to reach this potential, the dendrites nonlinearly converge to the balance potential. When the dendritic potential is below the separatrix, the compartment remains hyperpolarized. The voltage-dependent conductances open at lower input rates for longer dendrites, therefore they reach the E/I balance slightly earlier. The narrow range of inputs (<0.3 kHz) to reach the E/I balance in models with NMDARs implies that dendritic nonlinearities offer a mechanism for compensating electrotonic segregation in dendrites. In AMPA-only models, longer dendrites require additional 3 kHz of input to reach the E/I balance.

**Figure 2:**
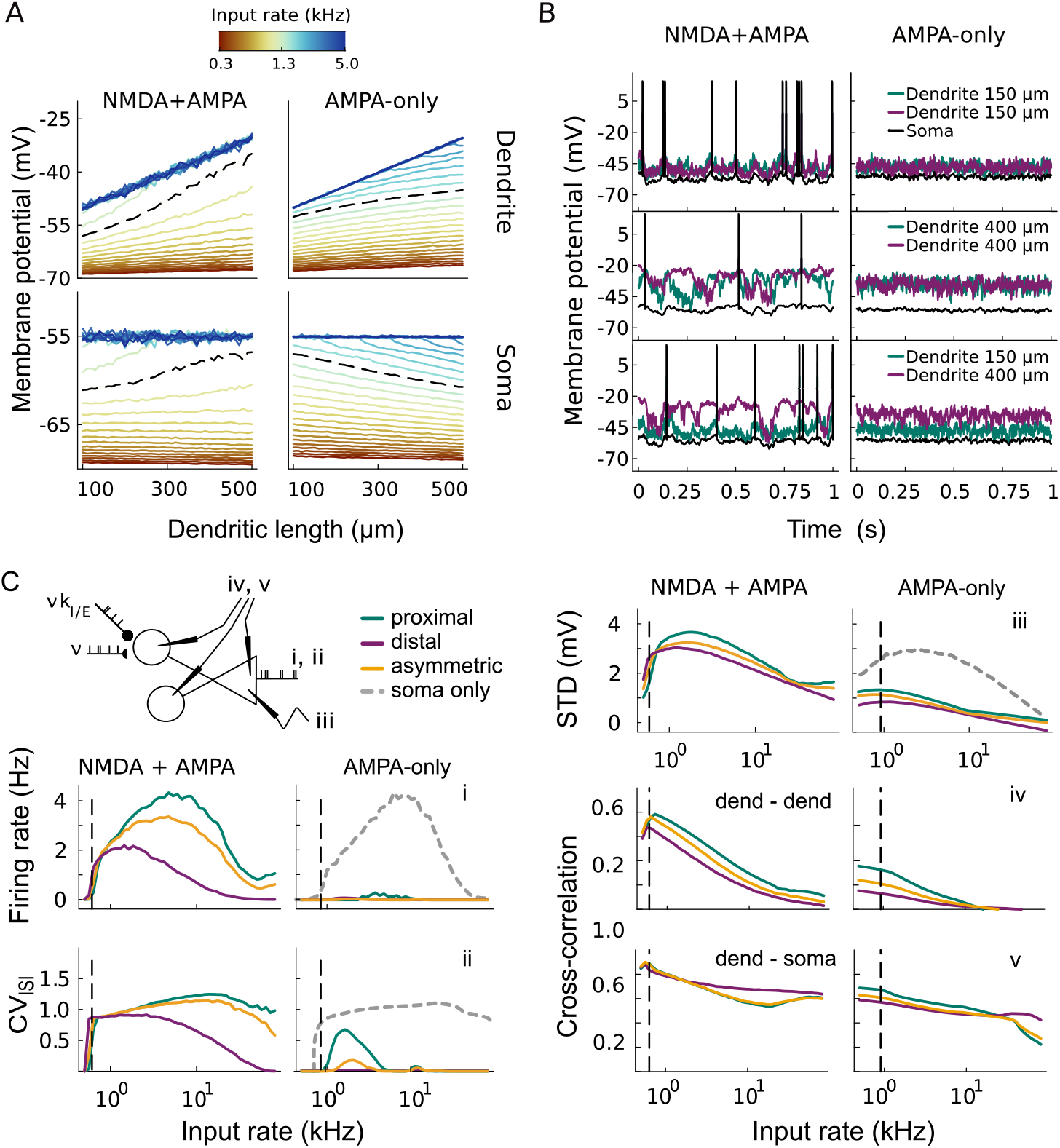
High conductance state in neurons with and without NMDARs. **A**, Average membrane potential of dendrites/soma for input rates from 0.3 kHz to 5 kHz (color code). Models with NMDARs reach the high-conductance state for input rates above 0.7 kHz (dashed line). AMPA-only models with short dendrites reach this state at 1.5 kHz (dashed line), but an increase to 3.5 kHz is required for long dendrites. **B**, Dynamics of membrane potential in dendrites and soma. Three dendritic configurations are shown, proximal-proximal (150–150 µm), distal-distal (400–400 µm), and proximal-distal (150–400 µm). For rigorous comparison, the same frozen spike train stimulates the compartments in all six simulations. Dendrites with NMDARs exhibit larger voltage fluctuations than AMPA-only ones and drive the soma to fire. **C**, Statistics of neuronal activity and membrane potentials in the balanced condition. 1600 neuronal configurations are averaged over three ranges of dendritic lengths: proximal (100 µm to 300 µm), distal (300 µm to 500 µm), and asymmetric (300 µm to 500 µm × 100 µm to 300 µm). For comparison, we show the soma-only model and parameters studied in Kuhn *et al*. (2004). Panels (**i**) and (**ii**) show the average somatic firing rate and its CV_ISI_. (**iii**) Standard deviation of the membrane potential in the free-membrane condition (no-spike threshold). (**iv**) Cross-correlation between the two dendritic branches and (**v**) between the dendrites and soma. Note that the dashed vertical lines in C correspond to the input rates identified in A.

**Figure 3:**
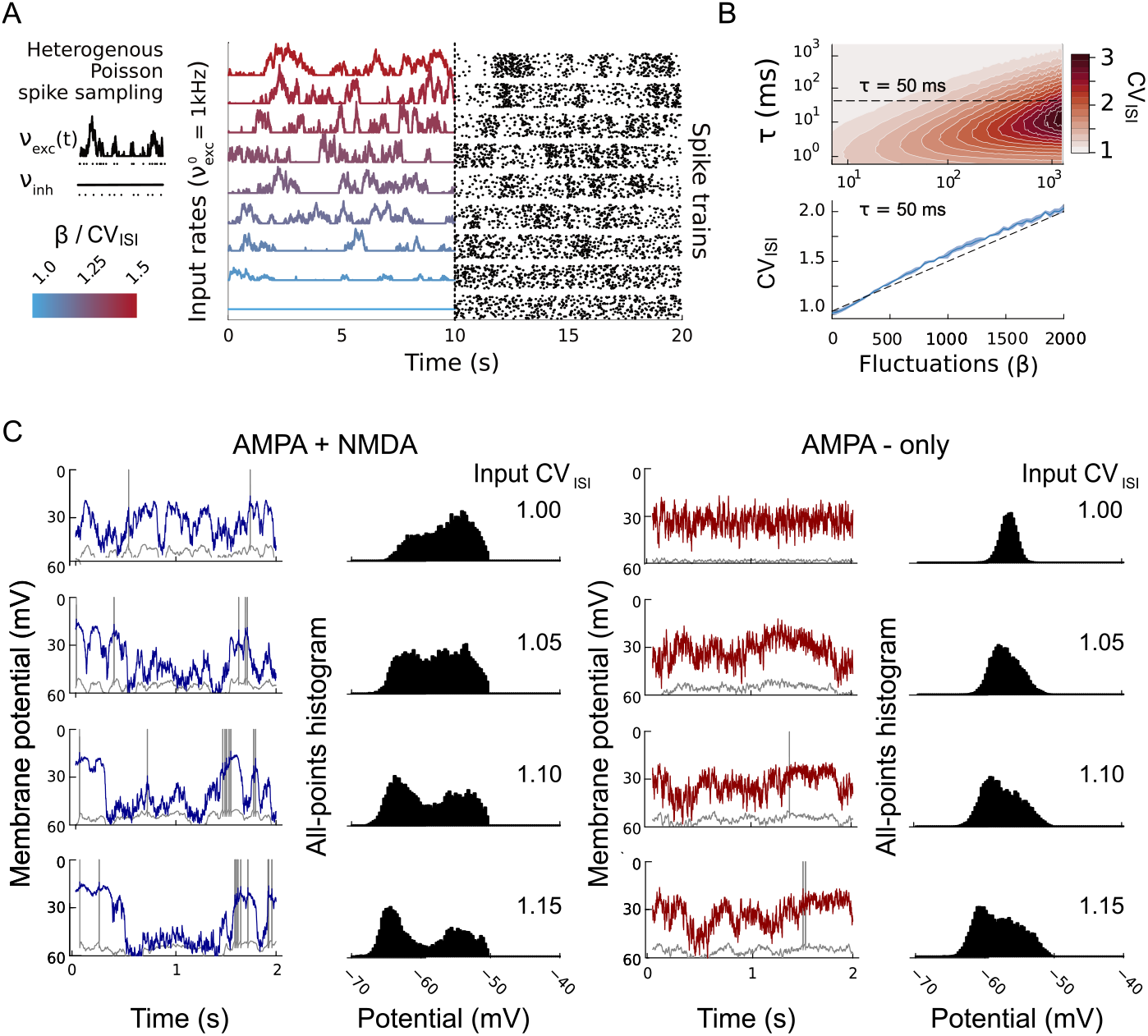
Input fluctuations lead to Up-Down states in models with NMDARs. **A**, Excitatory input rates and corresponding spike trains for increasing fluctuation sizes *β*. Input was drawn from an inhomogeneous Poisson distribution with a time-varying rate *ν*_*exc*_ (*t*) while *ν*_*inh*_ was held constant. Nine input rate samples are shown (left) with increasing CV_ISI_ (fluctuation size, *β*, blue to red), fifty spike trains sampled for each rate are shown on the right. With increasing *β*, signal fluctuations increase in magnitude, and spikes become less regular. Rates were scaled such that the average rate was fixed at 1 kHz. **B**, CV_ISI_ of the input spike train for *τ* = 0.1 ms to 1000 ms and *β* = 0 to 1000 on log-log scale. For *τ* =50 ms the CV_ISI_ of the spike train is linearly related to the parameter *β*. **C**, Membrane potential and APH of proximal-distal neuron, E/I-balanced at average input rate of 1.6 kHz with rate fluctuations *β* in the range of 1 to 1.15. With NMDARs, dendritic (blue) and somatic (grey) membrane potentials show large fluctuations and increasingly bimodal APH. Without NMDARs, the somatic membrane is less sensitive to *β*. Only one dendrite was plotted to facilitate comparison.

We also explored the neuron’s firing activity with and without NMDARs in different dendritic configurations. We used frozen noise input and applied the same excitatory and inhibitory spike trains to compare the simulations (Fig. 2*B*). The results indicate that the presence of NMDARs supports sparse firing activity and large fluctuations in the membrane potential, lasting in the order of 100 ms. Fluctuations on this scale are due to the self-regenerative processes linked to the voltage-dependent conductance in the NMDAR. When voltage-gated Ca^2+^-channels open, positive ionic currents flow through the membrane and drive a long-lasting depolarization, called NMDA-spike or plateau potential. This causes an upswing in the somatic potential and higher spike probability. In contrast, neurons with AMPA-only synapses undergo small, high-frequency fluctuations, which, combined with the low-pass filter integration in the soma, reduce the probability of somatic spiking. Thus, we observe that despite the model being in the same balanced condition, with the same average potential in soma and dendrites (dark blue in Fig. 2*A*), the physiology of NMDARs substantially changes the neuronal response.

To quantify our intuitions concerning the role of NMDA receptors, we measured the statistics of somatic firing (mean rate and CV_ISI_), the standard deviation of somatic membrane potentials, and the cross-correlations between the membrane potentials of different compartments for different input rates and dendritic configurations. For every input rate, only models with NMDARs exhibit somatic activity (Fig. 2*C* i) in the balanced condition, and its firing rate response is non-monotonic in the input rate. Neurons with distal dendrites show lower firing rates that reach their maximum at low input rates, whereas neurons with proximal dendrites fire more at higher frequencies. Asymmetric neurons display intermediate firing rates. Proximal and asymmetric neurons have comparable CV_ISI_ at approximately 1, and distal dendrites elicit a lower CV_ISI_, indicating more regular firing (Fig. 2*C* ii). Therefore, the somatic response depends on the geometry of the dendritic branch hosting the synapses. Cortical neurons, endowed with extensive and asymmetric dendritic arborization, can exhibit multiple regimes of synaptic integration, with long (short) branches being more responsive to low (high) input frequencies. Crucially, the results for the soma-only model (dashed line in Fig. 2C) are consistent with the results reported in (Kuhn *et al*., 2004), confirming the procedure used to obtain the E/I balance points.

The standard deviation of the somatic membrane potential shows fluctuations that are up to three times larger when NMDARs are present (compared to the AMPA-only condition), which explains the difference in firing rate across conditions Fig. 2*C* iii). Similarly, the cross-correlation between the membrane potentials of dendritic compartments is three times higher when NMDARs are present than with AMPA-only (Fig. 2*C* iv). Dendrites are indirectly coupled via the soma, hence their membrane potentials also correlate (Fig. 2*C* v), revealing that fluctuations in the dendrites are transmitted to the somatic compartment. High cross-correlation indicates that the entire neuron depolarizes and hyperpolarizes in unison, compensating for noise in the spike trains arriving at each dendrite and amplifying low-frequency fluctuations. As the firing rate increases and fluctuations become faster, the correlation between dendritic compartments reduces, and neurons enter a regime of branch-specific synaptic integration (low cross-correlations in iv and v of Fig. 1*C*). For low input regimes, the high cross-correlation between soma and dendrites suggests the presence of meta-stable states in the membrane dynamics. Soma and dendrites switch between depolarized and hyperpolarized states, reminiscent of the physiological Up-Down states.

### Dendritic non-linearity leads to Up-Down states under E/I input fluctuations

We have shown that under stationary E/I-balanced input, the membrane potentials of neurons with NMDARs correlate across compartments (Fig. 2*C* iv, v). *In vivo* network activity, however, is more variable (Paré *et al*., 1998; Maksimov *et al*., 2018) than a homogeneous Poisson process, and these network fluctuations have been associated with the occurrence of somatic Up-Down states (Wilson & Kawaguchi, 1996; Jercog *et al*., 2017; Papoutsi *et al*., 2014). To emulate network fluctuations we varied the CV_ISI_ of the input while keeping the input rate fixed (spike trains were implemented as inhomogeneous Poisson processes, details in Materials and Methods and Fig. 3*A* and Fig. 3*B*). Based on our observation that dendritic nonlinearities enhance the membrane’s sensitivity to correlations in the input, we expected that NMDARs support the occurrence of cellular UDS. We recorded the membrane potential of soma and dendrites for models with E/I-balanced input of 1.6 kHz and CV_ISI_ in the range of 1 to 1.15, with and without NMDARs (Fig. 3*C*).

When voltage-dependent NMDARs are present, the dendritic membrane potential shows excursions of more than 30 mV between the hyperpolarized and depolarized state, (Fig. 3*C*, left panels) whereas neurons without NMDARs were less sensitive to input fluctuations (Fig. 3*C*, right panels). Furthermore, somatic firing follows the fluctuations of the dendritic membrane potential. We used the all-points-histogram (APH, see Materials and Methods) to reveal the dwelling time of the membrane potential within each voltage interval. We observe that fluctuations in the input rate drive the soma of neurons with NMDARs into a bimodal regime. Without NMDARs, the soma has only one peak around − 60 mV. We quantified bimodality of the APH using Gaussian Kernel Density Estimation (GKDE, see Materials and Methods). With increasing window size, the number of peaks in the GKDE decreases (Fig. 4*A*). The bimodality index refers to the maximum kernel window that retains two peaks in the APH. We tracked the degree of bimodality in three dendritic models in different configurations (distal-distal, proximal-proximal, distal-proximal) and also tested a soma-only model without dendrites. Neurons with NMDARs exhibit high bimodality when the CV_ISI_ of the input exceeds 1.2 and input rates are low (0.5 kHz to 5 kHz, left panels in Fig. 4*B*), whereas it weakens at higher rates. This holds for both symmetric and asymmetric configurations when dendrites express NMDARs. In contrast, AMPA-only models (right panels in Fig. 4*B*) require substantial fluctuations to exhibit bimodality, and the index remains low. Interestingly, the soma-only model shows intermediate behavior, with a high bimodality index in a narrow range of input rates (4 kHz to 8 kHz), with and without NMDARs. Furthermore, fluctuations boost the firing activity in models with NMDARs, leading to bursting activity when at least one dendrite is short (Fig. 4*C*).

**Figure 4:**
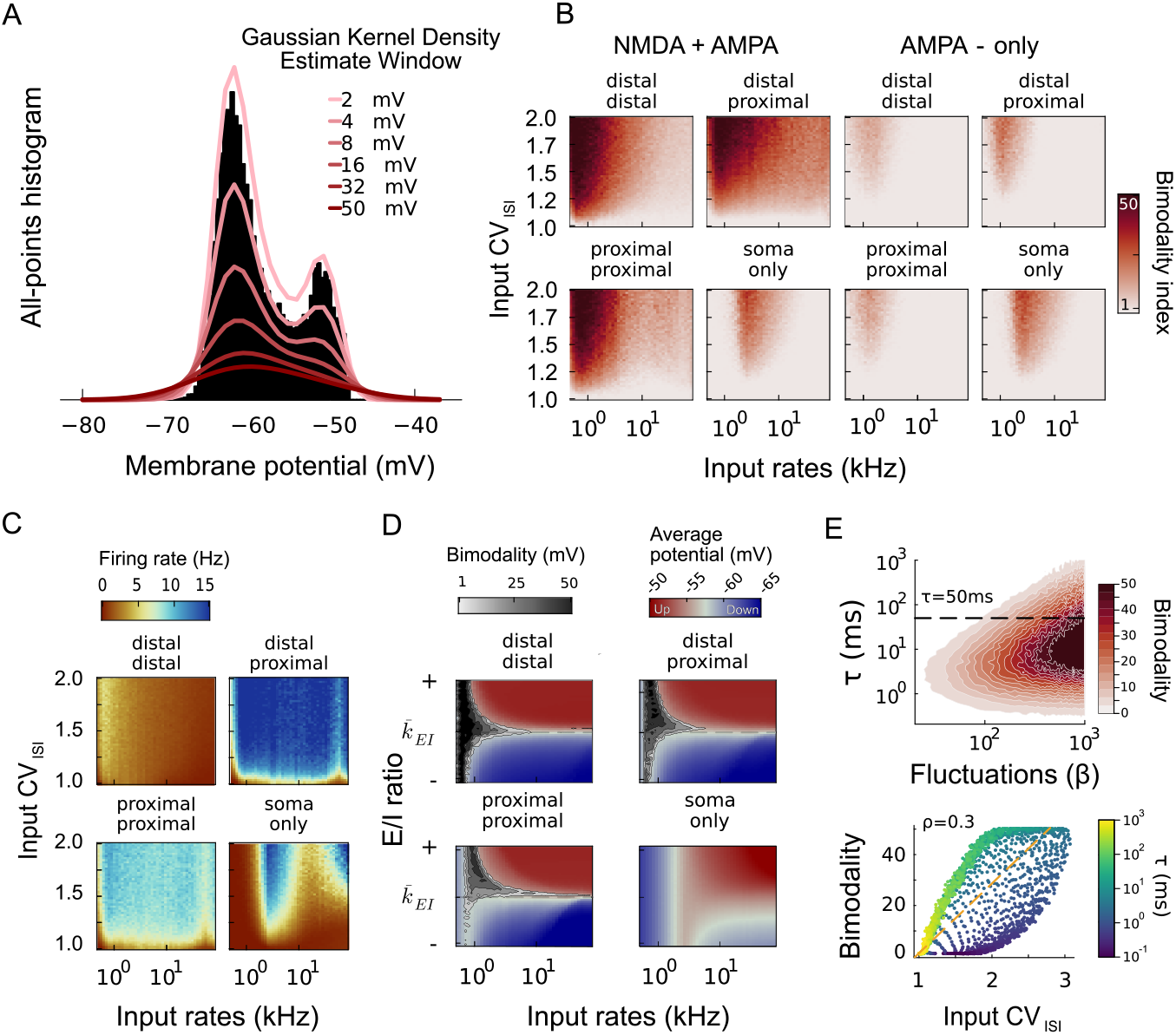
E/I balance and nonlinear dendritic integration support Up-Down states. **A**, APH for CV_ISI_ of 1.2, colored lines show GKDE for window sizes between 2 and 50 mV. The maximum window with two peaks in the GKDE defines the bimodality index. **B**, Bimodality index for different dendritic configurations and receptor composition, tested for input rates of 0.5 kHz to 80 kHz and increasing CV_ISI_. Neurons with NMDARs show stronger bimodality in the APH than AMPA-only neurons. **C**, Firing rates of models with NMDARs when input rate and CV_ISI_ are varied. For distal-distal, activity peaks at low input rates. With proximal dendrites, neurons burst during the Up-state. Firing rate increases sharply with CV_ISI_ across input rates. For soma-only, activity peaks at intermediate input rates regardless of fluctuation size. AMPA-only models (not shown) behave similarly. **D**, Bimodality (grey scale) and average membrane potential against variations in *k*_*E/I*_. For increasing input rates, the range of *k*_*E/I*_ for which the APH is bimodal shrinks around the E/I balance point 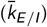. Outside the balanced state, the somatic membrane is permanently hyperpolarized (blue), or depolarized (red). **E**, Upper panel: bimodality of proximal-distal model for varying size *β* and timescale *τ* of fluctuations. Lower panel: bimodality increases with input CV_ISI_, regardless of fluctuation timescale (Pearson corr. *ρ* = 0.3). For *τ >* 10 ms, bimodality and CV_ISI_ correlate at *ρ* = 0.95.

We also examined the role of the E/I balance in eliciting Up-Down states. To this aim, we set the CV_ISI_ at 1.2 and measured the bimodality index while varying the E/I ratio (*k*_*E/I*_) in the range of 5 % to 200 % of the balanced 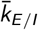. For input rates below 1 kHz, the E/I balance is not essential for the occurrence of UDS, but it becomes critical at higher input rates where the Up-Down states occur only within a narrow band of E/I ratios centered around 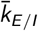 (Fig. 4*D*). The statistical properties of the input are crucial for the occurrence of UDS. We measured the bimodality index for varying fluctuation timescales *τ*_*r*_ and widths *β* (Eq. 13). Neither of them correlated with the bimodality index. However, bimodality was correlated with the CV_ISI_ of the generated spike train (Fig. 4*E*), i.e., the variability in the E/I-balanced barrage of afferent spikes drives the neuron into a bimodal state.

### Cellular and network origins of Up-Down states

We have so far identified dendritic nonlinearity as a crucial factor for somatic bimodality. Small fluctuations in the E/I-balanced state triggered UDS in models with NMDARs, whereas neurons with AMPA-only synapses required substantial excitatory input rate changes for UDS. Thus, bimodality arises from the interplay of cellular and local network dynamics, with NMDARs amplifying responses to cortical state fluctuations. However, Up-Down states have also been observed *in vivo* when the NMDAR antagonist MK-801 was administered (Smith *et al*., 2013; Palmer *et al*., 2014; Chen *et al*., 2012). We hypothesized that Up-Down states under MK-801 perfusion could be due to deep network oscillations in the anesthetized state of the recorded animals, where high input rate variability compensates for the role of NMDARs.

To test this hypothesis, we simulated a dendritic neuron model embedded in a medium-sized network (800 excitatory neurons, 200 interneurons) and recorded the membrane potential while gradually deactivating the cell’s voltage-dependent conductance (percentage of active NMDARs). Neurons received E/I-balanced inputs at 2 kHz, with CV_ISI_=1.2, and an oscillatory input with frequency of 1 Hz and amplitudes in the range of 0 kHz to 8 kHz, mimicking the cortical state under anesthesia (Fig. 5*A*). When the network was driven by sufficiently strong external modulation (4 kHz, or higher), the neuron entered the UDS dynamics regardless of the percentage of active NMDARs (Fig. 5*B*). When external modulation was weaker, only neurons with NMDARs active above 50% exhibited UDS (Fig. 5*C*, left). It also required 75% of active NMDARs for the neuron to fire. Direct comparison with a neuron receiving the same external fluctuations but no recurrent inputs (Fig. 5*C*, right) indicates that network reverberation was crucial in generating the neuronal Up-Down dynamics. Because the cells collectively switch between Up and Down states (Fig. 5*A*), the network nonlinearly amplifies the oscillatory input, alternating between high E/I firing (Up state) and inhibition-dominated activity (Down state). These input fluctuations are reflected in the recorded cell’s membrane dynamics and drive bimodality.

**Figure 5:**
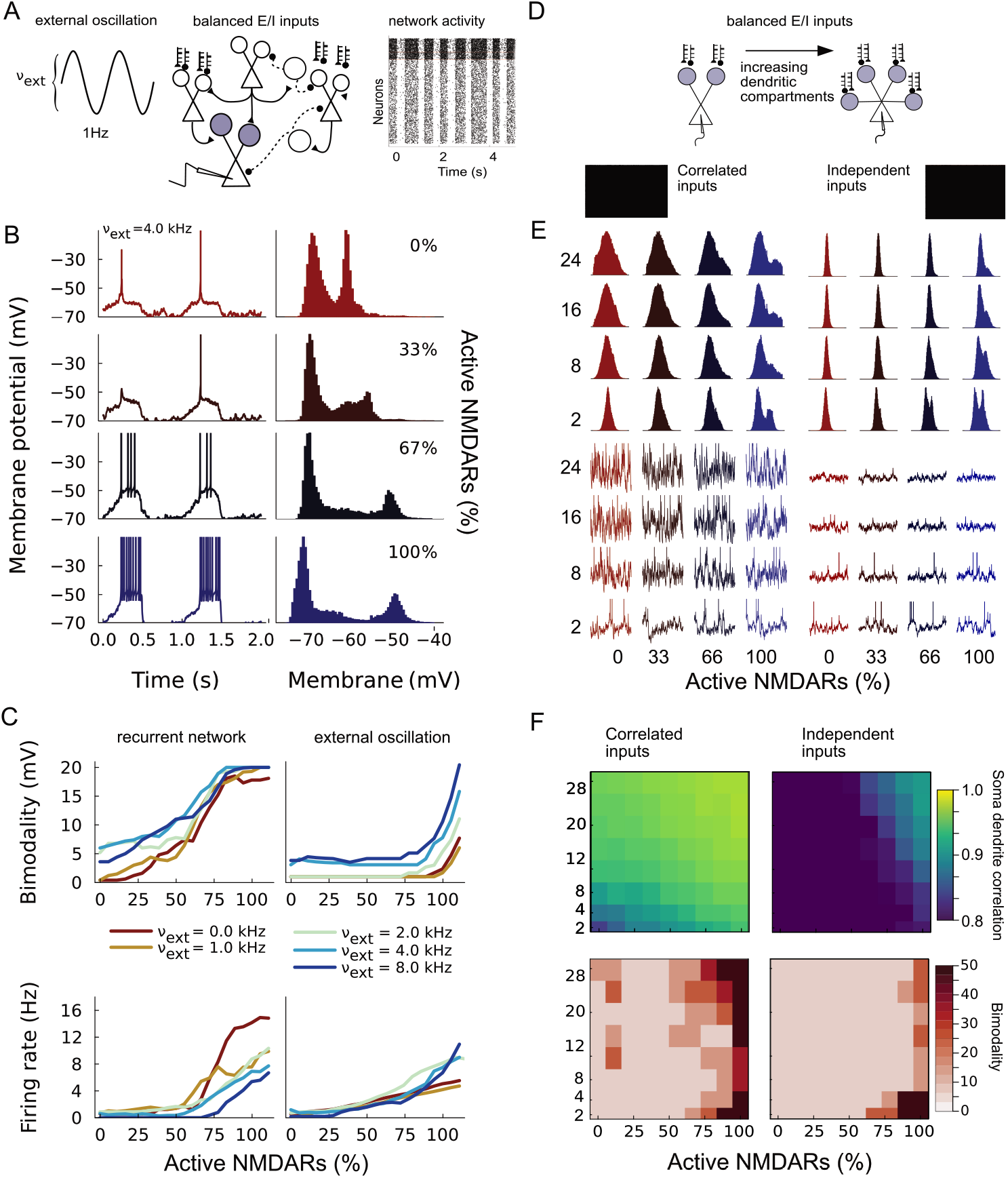
Network-induced UDS when NMDARs are blocked. **A**, The measured neuron (grey) receives excitatory and inhibitory afferents in a recurrent network. Network and measured neuron receive balanced input (2 kHz, CV_ISI_=1.2) along with a sinusoidal signal with frequency 1 Hz and variable amplitude *ν*_*ext*_. **B**, Somatic membrane potential and APHs for network-embedded cell and 4 kHz external input. Because of strong network oscillations, the neuron enters into the UDS regime when NMDARs are deactivated. **C**, Comparison of bimodality and firing rate in network-embedded vs. isolated neurons. Network amplifies the impact of external fluctuations and leads to bimodality when NMDARs are gradually deactivated. For isolated neurons, bimodality is observed only when NMDARs are near fully activated. **D**, Single neuron dynamics was investigated for varying number of dendritic compartments. Neurons received a 3 kHz signal (CV_ISI_= 1.2) with excitatory and inhibitory input spikes equally distributed over the dendritic tree. Excitatory spike trains are either correlated across dendrites or independent. **E**, APH and somatic membrane potential in the two input conditions for 2–24 dendritic compartments. For correlated inputs, neurons show large fluctuations in the membrane potential, regardless of the number of active NMDARs. For uncorrelated inputs, smaller fluctuations are observed. **F**, Cross-correlations between somatic and dendritic activity (top), and bimodality index for all tested neurons (bottom). Models with several dendritic branches show high cross-correlations but do not exhibit pronounced UDS. In these conditions, the high bimodality index reflects the bimodal hyperpolarized state shown in E, whereas in models with a smaller number of compartments, bimodality reflects UDS and is a direct consequence of active NMDARs.

We also tested whether the presence and combined effects of multiple dendritic compartments can enhance the amplification of input fluctuations. To this aim, we measured the UDS in neurons with 2–24 dendritic compartments receiving balanced inputs as above (Fig. 5*D*) while gradually deactivating NMDARs. We tested two conditions, one where input on the dendrites was correlated and one where it was independent (illustrated in Fig. 5*D*, left and right panels of Fig. 5*E, F*). When inputs were correlated, dendrites acted cooperatively, resulting in large somatic membrane fluctuations and high spiking activity, regardless of the number of compartments and the fraction of active NMDARs (Fig. 5*E, F*, left panels). In contrast, for uncorrelated inputs, increasing the number of dendritic compartments lead to a sharp and narrow distribution of somatic membrane potentials and largely abolished spiking activity and bimodality, regardless of the fraction of active NMDARs (Fig. 5*E, F*, right panels). Hence, multiple dendritic compartments can amplify the neuron’s responses to input fluctuations only when receiving correlated inputs.

Overall, these results show that Up-Down states can emerge in single cells even when voltage-dependent NMDA conductances are deactivated, if and only if an external, oscillatory network drive is sufficiently strong. These observations are consistent with experimentally observed UDS in anesthesized animals when NMDARs were partially blocked (Smith *et al*., 2013; Palmer *et al*., 2014; Sun *et al*., 2018) and demonstrate that, in those conditions, the emergence of Up-Down states is inherited from the network state which, under anesthesia, is more strongly synchronized. However, NMDARs allow single cells to detect and amplify fluctuations in asynchronous inputs, characteristic of the *awake* state, supporting the emergence of UDS even in the absence of network-level synchronization effects.

### Dynamics of NMDAR-driven Up-Down states

To determine whether the characteristics of Up-Down states that we observe in the model are compatible with physiological reports, we now characterize the dynamics of UDS in the neuron model and compare it to electrophysiological recordings *in vivo* from Anderson *et al*. (2000) and Wilson & Kawaguchi (1996). To reflect some of the heterogeneity in the dendritic tree of real pyramidal cells in our minimal model, we study it with two asymmetric (proximal-distal) dendritic compartments (150 µm–400 µm, respectively). The input firing rate and CV_ISI_ were based on the reference values for cortical activity proposed in Maksimov *et al*. (2018), with each dendrite receiving independent spike trains at 1.5 kHz and a CV_ISI_ of 1.2.

The presence of NMDARs allows the model to reproduce the characteristic UDS membrane dynamics observed in cortical recordings. However, the transitions occur at lower frequencies, as can be seen from a direct comparison (Fig. 6*A*) of the model activity and intracellular measures of complex cells in the primary visual cortex of anesthetized cats (Anderson *et al*., 2000). Notably, although stimulated with the same input spike trains, models with AMPARs only do not show marked UDS transitions. Following the experimental study, we quantified the membrane statistics and defined Up and Down states via the third and first quartiles of the distribution of membrane potentials. When the somatic membrane crosses the Up-state threshold (third quartile), a somatic Up state occurs which lasts until the potential crosses the Down threshold (first quartile). The average difference between the two states was 7.5 mV (p-value <0.001) for the NMDAR models and 3.0 mV (p-value <0.001) for the AMPA-only model, compared to experimental values in the range of 10 mV to 15 mV (Fig. 6B). The model with NMDARs also shows larger fluctuations in the Up state which is consistent with the higher power in the range of 20 Hz to 50 Hz that was reported experimentally. As observed in Fig. 3, larger fluctuations in the Up state drive the somatic activity which is consistent with the significant positive correlation between the instantaneous firing rate and the membrane potential (r=0.34, p-value <0.001) in the Up state that we observed only in the presence of NMDARs (Fig. 6*C*).

**Figure 6:**
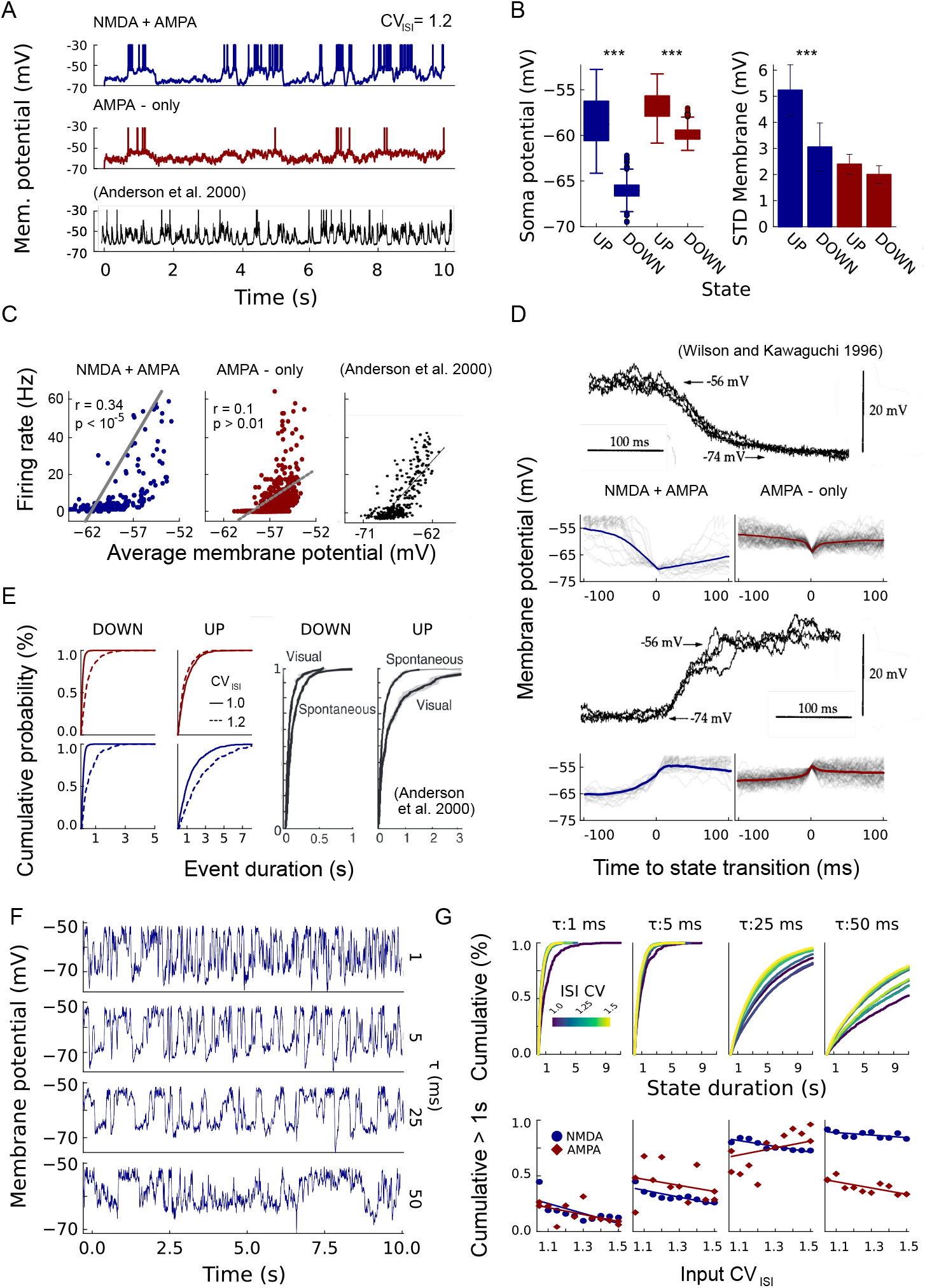
Models with NMDARs reproduce statistics of Up-Down states *in vivo*. **A**, Membrane potentials under fluctuating synaptic input (1.5 kHz and CV_ISI_ of 1.2) for dendritic models with/without NMDARs (blue/red), compared to cortical recordings in anesthetized cat (Anderson *et al*., 2000, black). **B**, UDS are identified as upper/lower quartiles of the distribution of somatic membrane states, collected over 10^3^ s. Average potentials of UDS differ significantly both with or without NMDARs but standard deviations in UDS differs significantly only in the presence of NMDARs. **C**, Average potential correlates significantly with instantaneous firing rate only in neurons with NMDARs. Experimental data in the right-most panel for comparison. **D**, Average membrane potential in proximity of UDS transitions, compared to *in vivo* measurements from Wilson & Kawaguchi (1996). With NMDARs, UDS transitions show an excursion between 10 mV to 20 mV and have a shape and time-course that more closely resembles the experimental data. For AMPA-only, transitions are faster and resemble statistical fluctuations rather than meta-stable states. **E**, Cumulative distribution of state duration for two input CV_ISI_, compared to the duration of UDS in anesthetized cat, with/without visual stimuli. In the models, average duration of Up states decreases with fluctuation size (red: AMPA-only, blue: AMPA+NMDA.). In the experiment, it increases in the presence of visual input. **F**, Membrane potential of asymmetric neuron with NMDARs for increasing fluctuation timescale (*τ* = 1 to 50). Longer timescales lower the intrinsic frequency of the UDS. **G**, Top: cumulative distribution of Up state duration for different fluctuation timescales. Color scale indicates the input CV_ISI_ in the range of 1.0 (dark blue) to 1.5 (yellow). Both *τ* and input CV_ISI_ affect the distribution of Up state duration. Bottom: probability of durations longer than 1 s from the cumulative distribution. Up-states are, on average, shorter for faster timescales and larger CV_ISI_.

The voltage-dependent receptors dominate the timescales of the transition between Up-Down states. In Fig. 6*D*, we averaged across all the transitions occurring in the 10^3^ s of simulation and compared the time course between the two states with recordings from Wilson & Kawaguchi (1996). In the presence of NMDARs, the onset and offset of Up states have a sigmoidal shape. The offset occurs in the range of 50 ms to 100 ms and it is slower than the onset (20 ms to 50 ms). Most importantly, the two states are visible before and after the transition. Without NMDARs, state transitions are less pronounced and faster than those observed *in vivo*. These results highlight the relevance of NMDAR-mediated events in the emergence of UDS whose core statistical properties closely resemble physiological evidence. However, they also suggest that there are important differences in the temporal structure, particularly in the frequency and duration of these events.

To further scrutinize these differences, we compare the temporal extent and distributions of UDS across models with experimental data of anesthetized animals in the presence or absence of visual inputs (data from Anderson *et al*., 2000). In this context, we hypothesized that visually evoked responses correspond to increased input correlations and larger CV_ISI_ (Karimipanah *et al*., 2017). Our model partially aligns with experimental observations concerning relative changes in the Up state in spontaneous versus evoked conditions (Fig. 6*E*). In both cases evoked activity (visual input) leads to longer, sustained Up-states, which are also observed in the model with NMDARs (dashed versus solid blue lines). However, the Up state duration was longer in the model and visual input did not lead to shorter Down states. Having full control over the timescale and amplitude of input fluctuations in our model, we systematically analyzed their impact on Up-state duration. We observed that lower CV_ISI_ and shorter autocorrelation timescales lead to faster, short-lived Up-states (*τ* = 1 ms to 50 ms, Fig. 6*F* and Fig. 6*G*). The best fit to the experimental data of Anderson *et al*. (2000), considering the modulation of input CV_ISI_ as a proxy for external stimuli, was obtained for input fluctuations with autocorrelation timescales on the order of 5 ms. This short timescale aligns with the fast responses of thalamic LGN neurons to drifting gratings (Siegle *et al*., 2021) thus supporting the idea that the modulation of Up-state lifetimes observed in Anderson *et al*. (2000) is a consequence of the interactions between thalamic inputs and dendritic NMDARs.

Taken together, the results presented in this section emphasize the significance of NMDAR-mediated effects in the origin of UDS in cortical cells. We conclude that voltage-dependent receptors enhance the emergence of marked UDS. However, the frequency and duration of these states mainly depend on the timescale of the input spike trains, that is, they are inherited from external inputs and recurrent network dynamics.

## Discussion

In this study, we investigated how dendritic nonlinearities mediated by voltage-dependent NMDA receptors shape neuronal responses to balanced excitatory/inhibitory synaptic bombardment. Using a *minimal*, three-compartment neuron model with two segregated dendritic elements, our results indicate that these nonlinearities detect and amplify input fluctuations and support the emergence of cortical Up-Down states (UDS). UDS have been observed in awake cortex, but are more pronounced in different stages of the sleep cycle and in anesthetized animals. The prevalent hypothesis is that changes in the neuronal state are driven by a combination of local network dynamics, glutamatergic inputs from the thalamus, and/or neuromodulation from the brain stem and basal forebrain (Tukker *et al*., 2020; Poulet & Crochet, 2019). In addition to external and intrinsic network drivers, cellular features such as electrogenic dendritic processes have been suggested to play a role in maintaining the Up state (Milojkovic *et al*., 2005; Antic *et al*., 2010), an observation that has been supported by theoretical and computational studies (Papoutsi *et al*., 2014). However, *in vivo* experiments have also shown that partial blockage of NMDARs does not seem to change the frequency of UDS transitions or the amplitude of the Up state depolarization (Palmer *et al*., 2014; Chen *et al*., 2013; Smith *et al*., 2013; Larkum, 2022), suggesting that while large, sustained depolarizations mediated by NMDARs can increase the spike-probability upon sensory stimulation, the UDS dynamics depends on a local balance of fast AMPA and GABA_A_ receptors throughout the dendritic tree.

The dendritic neuron model used here allowed us to elaborate on the role of dendritic processes and NMDAR-mediated nonlinearities by measuring the membrane dynamics when dendritic inputs were locally balanced. Cortical neurons recruit different mechanisms to reach and maintain a state of balanced excitation/inhibition (Turrigiano, 2011; Vogels *et al*., 2013; Hiratani & Fukai, 2017). We determined the E/I balance analytically and obtained a general solution for the ratio between excitatory and inhibitory firing rates that allow the cell to operate in a stable high-conductance state. We demonstrated that this ratio depends on dendritic geometry and the number of compartments, with longer dendrites requiring more excitation than shorter ones to establish somatic balance. These results are in agreement with empirical data on the distribution of excitatory and inhibitory dendritic spines (Iascone *et al*., 2020) and support the *dendritic democracy* hypothesis which posits the need for stronger synapses in distal regions (Häusser, 2001; Magee & Cook, 2000). Our study also suggests the (currently untested) hypothesis that input correlations at different timescales lead to different somatic responses depending on the dendritic location of clustered synaptic inputs. In particular, short dendrites should be sensitive to fast fluctuations whereas longer dendritic branches (e.g., the apical tuft) should be sensitive to fluctuations on longer timescales. This geometry-dependent response is a natural consequence of the integration timescale introduced by segregated dendritic compartments (Quaresima *et al*., 2022). The selectivity of dendritic branches with respect to input frequency is in line with recent physiological evidence that showed complex spatial arrangements of pre-synaptic axons onto distinct dendrites of sub-cortical and cortical cells (Callan *et al*., 2021; Lafourcade *et al*., 2022) and provides an elegant cellular basis for parallel processing and temporal multiplexing (Ujfalussy *et al*., 2018; Naud & Sprekeler, 2018).

Local, dendritic E/I balance can be achieved in our model regardless of the action of NMDARs, but these are strictly required for the cell to combine such local balance with supra-threshold somatic spiking and operate in physiologically realistic regimes. This indicates that AMPARs can coordinate local balance, but NMDARs are required for the cell to detect spatiotemporal coincidences in the input and elicit spike responses, in agreement with Larkum (2022). Passive segregated dendrites with AMPARs alone act as low-pass filters, attenuate fast fluctuations, and prevent spiking activity. Accordingly, isolated cells expressing only AMPARs require large external fluctuations to exhibit bimodality in the distribution of somatic membrane potentials. In these circumstances UDS dynamics is inherited from external inputs. Thus, our results suggest that the neuronal Up state and the maintenance of UDS reported under NMDAR blockage is the cellular-level consequence of network-level Up-Down dynamics (Compte *et al*., 2003; Holcman & Tsodyks, 2006), which is particularly prominent under anesthesia (Wilson & Kawaguchi, 1996; Destexhe *et al*., 2007). We tested this explanation by simulating neurons with gradually deactivated voltage-dependent NMDA conductances while being embedded in an E/I-balanced network with Up-Down dynamics. Despite partial, or even complete, deactivation of NMDARs, these cells showed bimodality in the all-points histogram, in agreement with experimental observations. Furthermore, we observed that the network Up state results in balanced E/I inputs to the neuron, whereas in the Down state, inhibitory inputs dominate, supporting previous proposals suggesting that Down states might be due to strong, dominant inhibition (Tartaglia & Brunel, 2017) and/or a transient absence of excitatory inputs (Dembrow & Spain, 2022).

While we aimed for a simplified neuron model that could integrate sub-cellular and network-level effects and retain physiological consistency, it is possible that this simplicity limits the scope of our conclusions. The computational findings by Benucci *et al*. (2004) suggest that the temporal filtering introduced by many dendritic cables with heterogeneous timescales supports the somatic Up state via the generation of broad and continuous depolarizing dendritic currents. In agreement with this hypothesis, we observed that increasing the number of dendritic compartments increases dendritic cross-correlations, and even more so in models equipped with NMDARs. Thus, complex dendritic arborizations and dendritic NMDARs cooperate in increasing the neuron’s sensitivity to external fluctuations. However, inputs on different branches are uncorrelated, the bimodality index decreases for models with a larger number of dendritic compartments. This observation suggests that NMDAR-mediated Up-states require the concurrent depolarization of multiple dendritic compartments. In our simplified model where each dendritic compartment corresponds to a large section of the neuron’s arborization, these cooperative effects are easier to instantiate, whereas more detailed morphological descriptions require a more careful control over the inputs driving each individual branch. In any case, local cooperativity due to branch- or compartment-specific input clustering has been proposed as a fundamental computational advantage of dendrites in cortical neurons (Ujfalussy *et al*., 2018). Regardless of the number of compartments considered, we showed that NMDA receptors boost coordinated fluctuations in the cell, resulting in high (*>*0.5) cross-correlations among compartments. These results align with whole-cell recordings in pyramidal neurons (Waters & Helmchen, 2004), account for the correlations between local and global events observed in single branches *in vivo* (Otor *et al*., 2022; Russell *et al*., 2022; Palmer *et al*., 2014), and indicate that electrical information propagates more efficiently in nonlinear dendrites.

Finally, when comparing the statistical properties of UDS in our model with those observed experimentally, we showed that the presence of NMDARs yields activity patterns that are consistent with electrophysiological recordings from Anderson *et al*. (2000) and exhibit similar state transition profiles (Up-Down and Down-Up transitions) as in Wilson & Kawaguchi (1996). To capture the relative distribution of state durations in ongoing versus evoked conditions (mirroring the results reported in Anderson *et al*., 2000), we show that the timescale of input fluctuations is critical and well-aligned with intrinsic timescales of LGN neurons upon visual stimulation (Siegle *et al*., 2021).

This study shows that dendritic nonlinearities mediated by NMDARs can explain the emergence and maintenance of UDS under realistic *in vivo*-like conditions. Their effects can selectively detect and amplify input fluctuations even in the presence of asynchronous inputs. While the UDS can be passively inherited from external inputs (intrinsic or extrinsic oscillations), NMDAR-mediated dendritic nonlinearities are required to yield robust, physiologically compatible responses. Importantly, the temporal features of Up- and Down-states (duration and switching frequency) appear to be dictated by the characteristics of external modulations (timescale and size).

## Notes

### Competing Interest Statement

The authors have declared no competing interest.

### Summary of Updates

The article has been revised in the following parts: 1. Analytical derivation of the balance condition in dendrites. 2. Up Down activity under network oscillations 3. Up Down states in neurons with multiple (N>2)compartments. 4. Removal of Supplementary Materials

